# Comparing frontal inclination angles measured from digital 3D models and 2D photographs of dry male and female crania from Croatia

**DOI:** 10.1101/2021.01.04.425183

**Authors:** Anja Petaros, Sabrina Sholts, Mislav Čavka, Mario Slaus, Sebastian K.T.S. Wärmländer

## Abstract

The frontal bone is one of the sexually dimorphic elements of the human skull that can be used for sex estimation of unidentified human remains. Numerous morphological features of the frontal bone, such as its angle of inclination, maximum anterior projection (glabella), and rounded elevations (frontal eminences) have been shown to differ between males and females. Various approaches have been developed to assess the frontal inclination in particular, and recently a method has been proposed where the angle of the frontal slope is measured from snapshots of digital three-dimensional (3D) models of human crania. However, as 3D-based investigations of skeletal material can be time-consuming and expensive, we here compare measurements of frontal angle inclination from 3D model snapshots to measurements from 2D photographs for a large sample (61 females and 61 males) of dry archaeological crania from medieval Croatia. Although angles measured from 3D snapshots and 2D photographs produced discriminant functions that classified crania by sex with similar accuracy (around 73%), the angles recorded from the 2D photographs were systematically one degree smaller than the angles recorded from the 3D images. Thus, even though both data sets were useful for sex estimation, we conclude that angles measured with the two different techniques should not be combined.

## 1. INTRODUCTION

Our perception of a head as feminine or masculine is mostly related to sexually dimorphic traits located in the upper third of the face [1], such as the anterior (glabellar) protrusion, the dimensions of the frontal eminences and sinuses, and the overall shape of the frontal bone. The development of these traits is influenced by numerous factors, including sex differences in maturation time, growth vectors, and the separation of the inner and outer tables of the frontal bone. The resulting shape dimorphism makes the frontal bone a valuable resource for sex estimation from the cranium, as it is well established that the slope of the frontal bone is larger in males than in females [2–5].

Sex estimation of the adult skeleton is a fundamental step in constructing the biological profile of an unknown individual, in both archaeological and forensic contexts [6, 7]. For incomplete skeletons, sex estimations are often based on the skull [2, 8]. The male skull is generally larger than the female, but adult human skulls exhibit sex differences in dozens of morphological features, which together are more diagnostic than size alone [9].

Few textbooks however recommend using the inclination angle when sexing unidentified remains, mainly because the large observer error associated with visual inspection of the forehead when it is assessed with simple descriptors (i.e. sloped/vertical) [10, 11] or ordinal scales, e.g. from −2 to 2 [12]. Craniometric measurements of the frontal bone have proven more successful [13, 14], and a method where the angle of the frontal slope is measured from digital three-dimensional (3D) models of crania has recently been proposed [5].

The frontal angle is a continuous parameter that can be measured with little observer error and which is easily incorporated into statistical analysis, making it well-suited for modern forensic analysis [10–12, 15–20]. Although recent advances in three-dimensional (3D) imaging technology and digital morphometric analysis have allowed researchers to quantify cranial features with advanced statistical analyses [21–23], 3D-based investigations of skeletal material can be time-consuming and expensive. Thus, in this study we investigate if the previously proposed method for measuring the frontal inclination from digital 3D models [5] can be used with 2D photographs as the data source.

Our sample consists of a large number (n=122) of dry archaeological crania from mediaeval Croatia, which were photographed in standard lateral position. 3D models were created with a CT scanner, and captured with 2D screenshots. Both types of images were then used to record trait scores as well as frontal inclination angles defined by the osteometric points glabella and supraglabella. Next, discriminant functions for sex estimation were created based on the different data, which allowed us to compare the usefulness of 2D photographs and 3D models for measuring frontal inclination angles for sex estimation.

## 2. MATERIALS AND METHODS

### Sample

The sample for this study consists of 122 human crania (61 females and 61 males) from the osteological collection of the Croatian Academy of Arts and Sciences (Zagreb, Croatia). All crania belong to adult individuals (determined by the fusion of long bones and eruption of third molar) with well preserved and complete skeletons and no antemortem head trauma. Sex of the skeletons was assessed using standard anthropological methods of visual inspection of the pelvis together with discriminant function analysis of long bone length, applying existing formulae developed for the medieval Croatian population [24, 25]. The skeletons have been excavated from four archaeological sites in the Dalmatian region of southern Croatia, and are dated to the Early (9^th^-11^th^ c.) and Late (12^th^-16^th^ c.) Croatian Medieval Periods (Table 1).

**Table 1.**
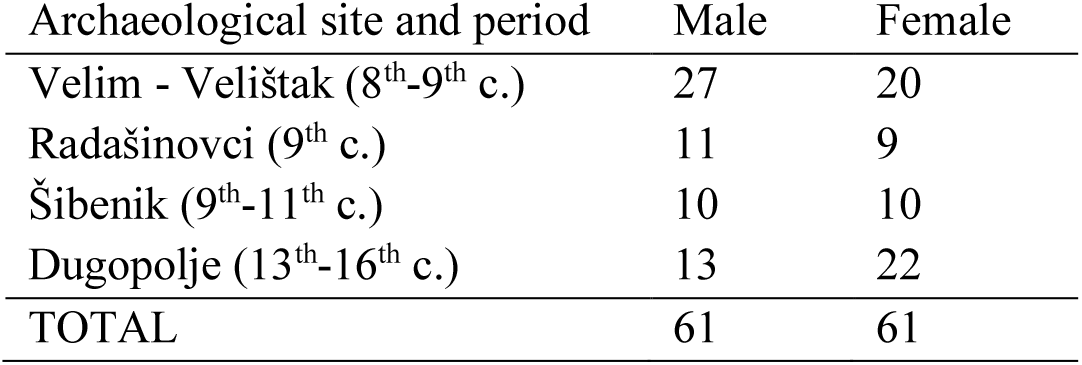
Demographic characteristics of the sample.

### Photography and 3D imaging

2D colour photographs of the crania positioned in left lateral profile (Fig. 1a) were recorded with a digital camera (SONY Cybershot DSC-HX300). In order to standardize the position of each cranium, the FOROST Cranial Photography Protocol [26] was used and adapted to the limited photographing material available at the collection spot, e.g., since a copy stand was not available, the camera was fixed on a shelf.

**Figure 1.**
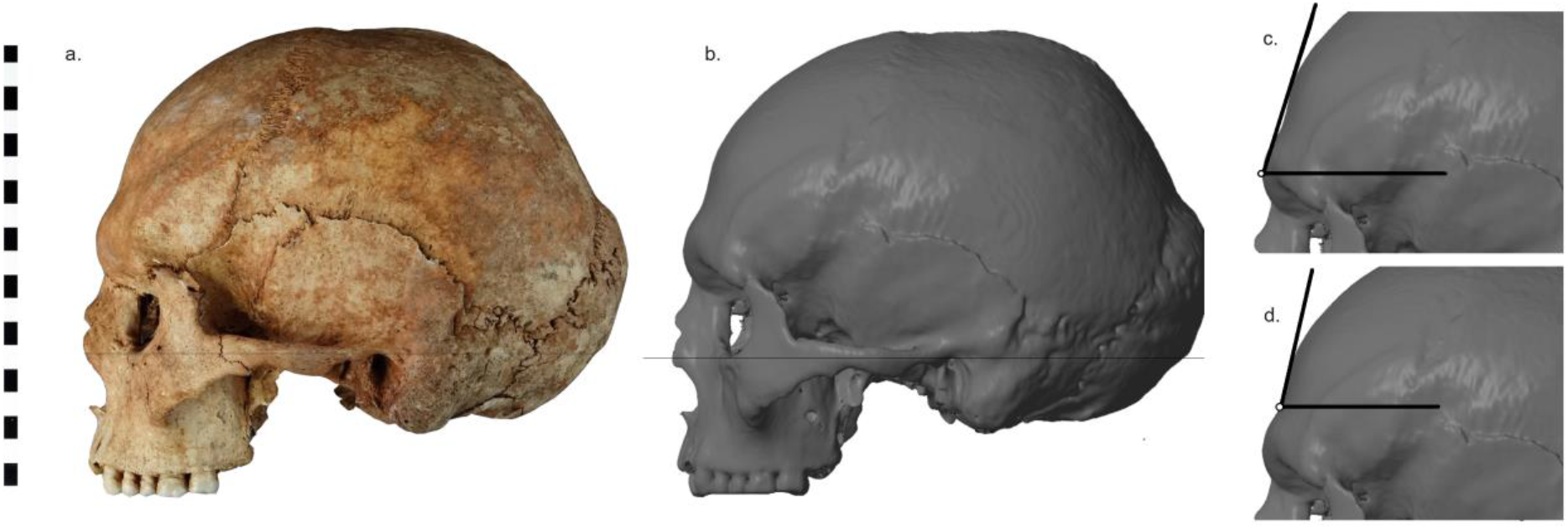
2D photograph (a) and 3D model snapshot (b) of a cranium, showing the orientation used for the inclination measurements. The frontal inclination angles were measured either from glabella (c) or supraglabella (d), in relation to the Frankfort horizontal plane. The scale to the left is in units of centimeters.

For 3D imaging, the crania were scanned at the Department of Diagnostic and Interventional Radiology, University Hospital Dubrava, Zagreb, Croatia, using a MDCT unit (Sensation 16, Siemens AG Medical Solutions, Erlangen, Germany) operating at 120 kV/320 mA and recording continuous layers without overlap, using 12 × 0.75 mm collimation. The resulting DICOM data (approximately 250 slices per cranium) were imported into the 3D Doctor imaging program (Able Software Corp., 1998-2011). A neutral tissue kernel was used for CT image reconstruction, followed by threshold-based bone extraction. The lower and upper threshold values were defined semi-automatically. After segmentation was completed, all crania were reconstructed using surface-rendering, and exported as stereolitography (STL) models. These 3D models were then oriented along the Frankfurt Horizontal (FH) plane and adjusted for left-right symmetry using the free version of the Netfabb Studio software (netfabb GmbH, Germany, 2009), after which a 2D screenshot of the left lateral profile was recorded (Fig. 1.b).

### Inclination measurement

All lateral profile images (i.e. both photographs and 3D model screenshots) were checked for orientation along the FH plane: where deviation was noted, the images were rotated digitally and adjusted to get a proper alignment of the upper auditory meatus and the inferior orbital rim (Fig 1). Using the 3D Doctor software, which allows automated angle measurements, frontal inclination angles were recorded using two lines intersecting at glabella: 1) a line parallel to the FH at the most prominent point of the glabellar region (effectively defined as the most prominent upper pixel in the region) and 2) a line tangent to the frontal bone outline at glabella (Fig. 2c). In addition, the inclination angle from supraglabella (defined as the point of maximum concavity behind the glabella in the midline) was also measured (Fig. 2d)). To evaluate observer error, blind measurements of the inclination angles were performed by one observer (author AP) over multiple days and with images presented in a randomized order.

**Figure 2.**
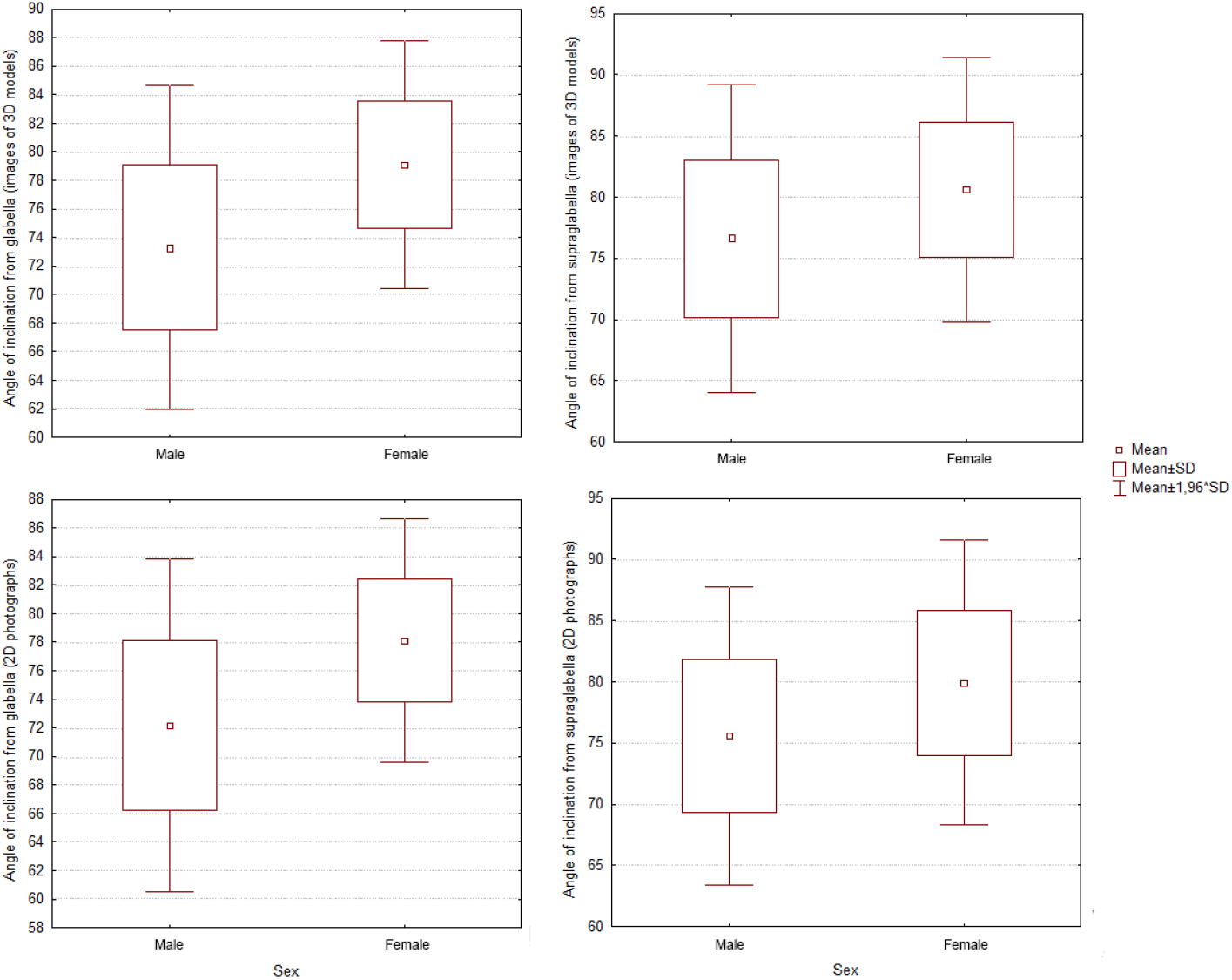
Distribution of inclination angles for male and female crania, measured either from 2D photographs or from 3D models.

### Trait scoring

Using the lateral profile images, crania were visually assessed for frontal inclination type, using the vertical-rounded-full (i) or low-sloped (ii) categories described by Rogers [10]. From the same images, the glabellar eminence was scored as a single expression from one to five, following descriptions by Buikstra and Ubelaker [27]. Repeated blind scoring trials by two observers (authors AP and SBS) were performed over multiple days, with the images presented in a randomized order with their ID numbers blinded.

## 3. RESULTS

### Inclination angles

The results of the inclination angle measurements are presented in Table 2 and Fig. 2. The complete data sets are presented in the supplementary material. It is clear that males display smaller inclination angles than females, and that angles measured from 3D model screenshots are larger than angles measured from 2D photographs. For inclination angles measured from glabella, the average male/female values are respectively 73.3° and 79.1° when measured from the 3D models, and 72.2° and 78.1° when measured from the 2D photographs (Table 2). For inclination angles measured from supraglabella, the average male/female values are respectively 76.6° and 80.6° when measured from the 3D models, and 75.6° and 79.9° when measured from the 2D photographs (Table 2). Thus, the difference between angles measured from 3D models and 2D photographs are around 1°, and paired t-tests show that these differences are significant at p<0.05 for all four data sets (Table 2). The male/female difference is around 5.8° for glabellar angles, and 4.2° for supraglabellar angles. These differences are highly significant at p<0.0005 for all four data sets (Table 2).

**Table 2.**
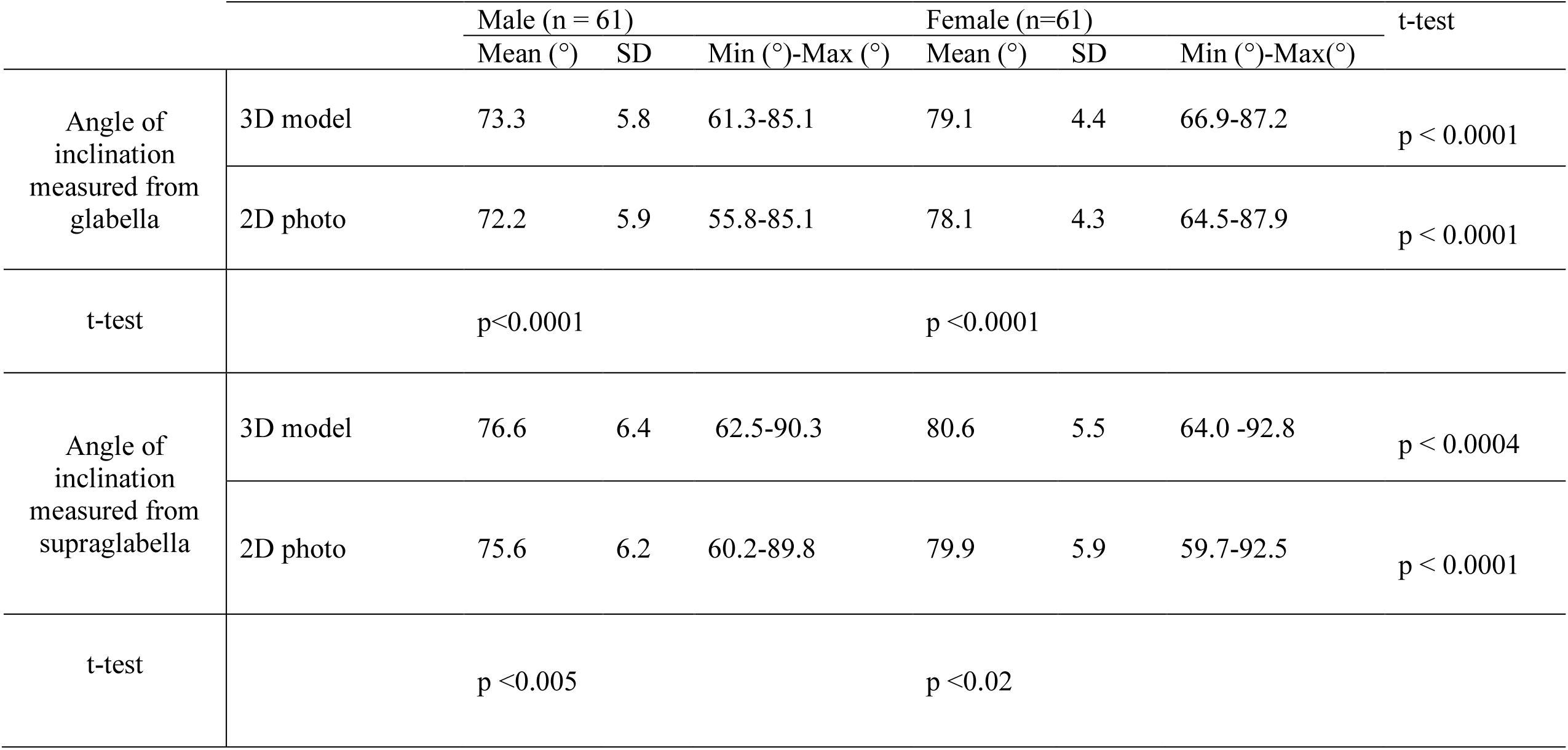
Angles of inclination measured from both glabella and supraglabella, for male and female crania. Data were obtained from 3D model screenshots and from 2D photographs. Double-sided t-tests for independent samples were used to evaluate the differences between male and female inclinations, while paired (dependent) double-sided t-tests were used to evaluate the differences between angles recorded from 3D model screenshots and 2D photographs.

Because of the larger difference for glabellar angles, we expected these data to perform better in statistical sex estimations. Furthermore, as the female inclination angles are less variable than the male ones, we expected statistical estimations to perform better for female crania. These expectations were largely met when discriminant functions for sex determination were calculated based on the inclination angles (using the Statistica 8 software from StatSoft, Inc., USA, 1984-2006). The results, presented in Table 3, show that discriminant functions based on glabellar inclination angles performed better (around 73% correct classifications) than predictions based on supraglabellar angles (slightly above 60% correct classifications). The predictions were more accurate for females (77% using glabellar angles) than for males (around 70% using glabellar angles).

**Table 3.** Number of correct sex predictions from discriminant function analysis, where the inclination angles of the male and female crania were measured either from glabella or from supraglabella, from 3D models and from 2D photographs.

| Inclination angle | Sex | 3D model | 2D photo |
| --- | --- | --- | --- |
| glabella | Male | 43/61 (70.5%) | 42/61 (68.9%) |
|  | Female | 47/61 (77.0%) | 47/61 (77.0%) |
|  | All | **90/122 (73.8%)** | **89/122 (73.0%)** |
| supraglabella | Male | 37/61 (60.7%) | 38/61 (62.3%) |
|  | Female | 37/61 (60.7%) | 40/61 (65.6%) |
|  | All | **74/122 (60.7%)** | **78/122 (63.9%)** |

### Measurement errors for the inclination angles

All measured inclination angles are presented in the supplementary material. The mean difference between the first and second measurements by observer AP for the glabellar angle recorded from the 3D models was 0.20° (range 0.00° to 0.65°), which was less than the 0.42° for the supraglabellar angle (range 0.00° to 3.14°). For measurements from the 2D photographs, the average variation was higher: 0.31° (0.01°-0.95°) for the glabellar and 0.53° (0.0°-1.85°) for the supraglabellar angle. Thus, the glabellar angle appears to be more reproducible than the supraglabellar angle (both for 2D and 3D data), and the images from the 3D models appear to yield less observer variation in the angle measurements than the 2D photographs.

### Trait scores

Trait scores for glabellar prominence (ranked 1-5) and frontal inclination (sloped/vertical) were evaluated by two observers (AP and SBS). The results are shown in Table 4, while complete data sets are presented in the supplemental material. As expected, male crania yield higher glabella scores (mean value around 3.5) and a larger proportion of “sloped” frontal inclinations (69% when measured from 3D model data, 82% when measured from 2D photographs) than female crania (mean glabella score around 2.2; 31% sloped frontal inclinations measured from 3D model data, 41% from 2D photographs). Discriminant functions based on these data classify sex with good accuracies: around 80% correct classifications with the glabella trait scores, and 70% using sloped/vertical inclination. These results are slightly better than the predictions based on inclination angles, and thus clearly demonstrate the usefulness of trait scoring for sex determination, even though the trait-based sex predictions performed better for males than for females.

**Table 4.**
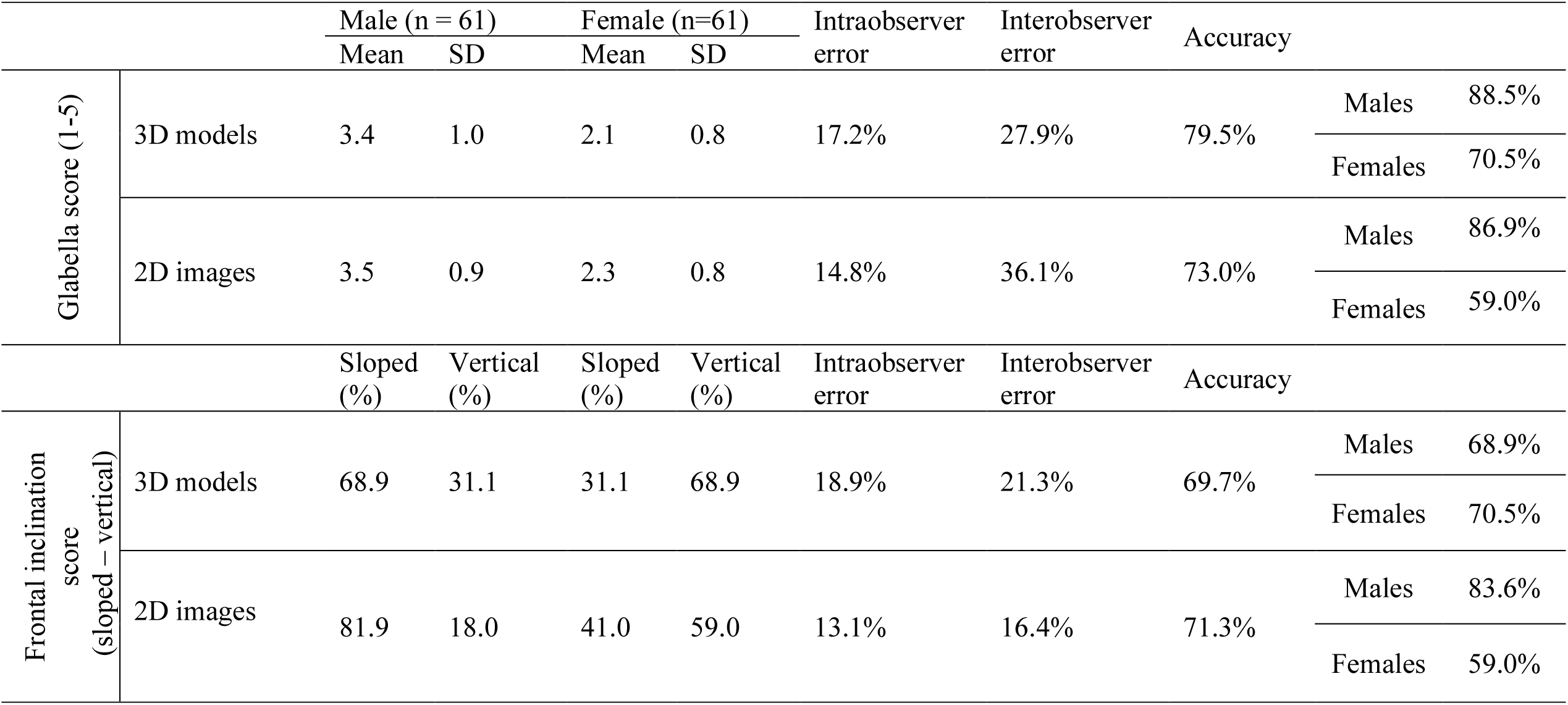
Glabella and Frontal inclination trait scores for male and female crania, derived from visual analysis of both 3D models and 2D photographs, presented together with observer errors and accuracy values derived from discriminant functions analysis. The observer errors were calculated as the proportion of crania that were given different scores between two trials. The presented results were calculated from two scoring trials conducted by author AP, except for the interobserver error which was calculated from one scoring trial by author AP and one scoring trial by author SBS.

Interobserver errors for the trait scores were in the range 16% – 36%, while intraobserver errors were never greater than 20% with lower for scores obtained from 2D photographs than from 3D models (Table 4). Thus, the observer error rates were similar to the rate of misclassified crania. Inclination scores from 3D model data and from 2D images were found to differ in 20 of the 122 specimens (i.e. 16.4%), similar to the observer errors. Glabella trait scores obtained from 3D model data and from 2D images, however, differed for 39.3% of the sample.

### Correlation between glabellar prominence and inclination angles

Spearman’s rank correlation coefficients (SRCCs) were calculated to investigate the dependence between the protrusion of the glabella (scores 1-5) and the angles of inclination (continuous parameter) measured from either glabella or supraglabella. For angles and scores taken from the 3D models, the correlation was 0.64 for the inclination angles from glabella and 0.43 for the angles from supraglabella. For data from the 2D photographs, the correlation was 0.62 for glabella inclination angles and 0.39 for supraglabella angles.

## DISCUSSION

It has previously been shown that frontal inclination can be successfully measured from digital 3D models [5], and our current results are in line with those observations. Measured from glabella using 3D models, the average male and female frontal inclination angles for the Croatian sample (73.3° and 79.1°; Table 2) are similar to those for European and American whites, but smaller than for American blacks and Chinese [5]. The supraglabellar inclination angles display smaller differences between populations [5], making comparisons difficult. While a detailed analysis of Croatian cranial shape still has to be carried out, the current results suggest that at least the frontal bone of the Croatian sample appears to display typical European features. So far, detailed investigations of frontal bone shape have only been carried out for a handful of populations. A more posterior position of the frontotemporale was observed in American White males [28], and a more protruded glabella together with a flattened and sloped frontal region was found in a sample of Central European males [29]. For African remains, the profile of the forehead was found to be the second-best feature for sex determination [30], while modern Japanese crania showed no sex differences in frontal bone inclination when analysed with semi-landmarks and Bezier curves [31].

The current inclination angles were measured both from 2D photographs and from screenshots of 3D models with high precision, i.e. ±0.2 degrees between repeats for the glabellar angles and ±0.5 degrees for the supraglabellar angles (Supplementary Table S2). Interestingly, the inclination angles measured from 2D photographs are about 1° smaller than the corresponding angles derived from 3D models, for both glabellar and supraglabellar angles and for both male and female crania (Table 2). Although small, the difference is very consistent and statistically highly significant (Table 2). There are several possible reasons for this discrepancy: First, the tools present in 3D software – such as digital rulers and automated symmetry alignment – make it easier to align 3D models along the FH plane, compared to physical crania. This is especially true for damaged crania, where the physical specimens sometimes tend to tilt to one side, affecting FH alignment and thus the inclination angle measurements. Second, the low-resolution screenshots for the 3D models suffer from pixilation, with the drawback that shape information is lost when the frontal profile is not displayed as a smooth curve. The advantage, for inclination measurements, is that it becomes relatively easy to repeatedly identify the “most prominent upper pixel” in the glabellar region. For the high-resolution photographs, the smooth curve of the glabellar region arguably makes it more difficult to repeatedly locate the same starting point for the inclination angle. Thus, the pixilation effect improves the repeatability (precision) but not necessarily the accuracy of the measurements. This might explain why angles measured from 3D model screenshots showed smaller intraobserver errors (variation) than angles measured from 2D photographs (Supplementary Table S2). Third, the cranial profile captured with the photograph will vary slightly depending on the optical properties of the camera lens as well as the geometry (distance and angle) between the camera lens and the object. This variation in projection will influence both the measurement of the frontal angle and any attempts to define the FH plane from cranial landmarks identified in the photograph. For 3D models, the 3D software may map the 3D object onto the 2D screen either in orthographic (orthogonal) projection or in perspective projection, but the observer can easily switch between different projections (we used orthographic projection). Differences in projection lead to mismatch in frontal profiles when 2D photographs and 3D image screenshots of the same crania are superimposed (Fig. 3), and this is the most likely reason for the systematic difference between the angles derived from the 2D and 3D images. The effects of image projection on craniometric measurements clearly deserve to be further investigated in a separate study.

**Figure 3.**
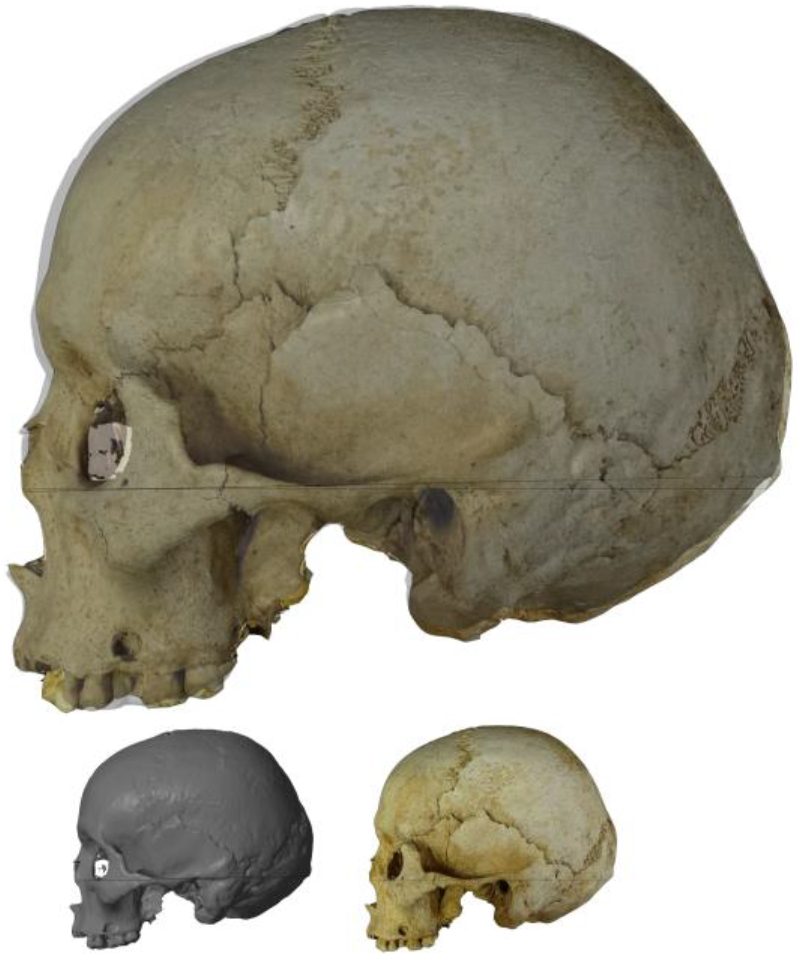
Overlaid 2D photo and 3D model snapshot of cranium R42, displaying clear differences between the two images.

The consistently smaller inclination angles measured from the photographs show that such data should never be combined with angles measured from 3D models (and vice versa). The discrepancy raises also the question whether one type of image should be preferred over the other. Although creating 3D models can be both laborious and expensive, producing screenshots of such models is easier than recording 2D photographs of physical crania, especially if a database with 3D models already is available. Aligning a 3D model in a desired orientation is straightforward, while the orientation of the 2D photograph is fixed and its perspective projection depends on the camera setup and the camera lens properties. Because these parameters typically are not documented, comparisons between photography-based studies are prone to have inherent sources of error. Osteometric measurements from 3D models may vary with the resolution of the model [32], but such resolution effects will likely become smaller as 3D software algorithms and 3D scanner resolution are improved, and as storage media for large 3D files become more affordable. Thus, we believe that 3D-based approaches are better suited for use in applied forensics.

Statistically significant (p < 0.001) difference in frontal inclination between the sexes were observed for both glabellar and supraglabellar angles, for measurement from 2D photographs as well as from 3D model screenshots (Table 2). Discriminant functions based on glabellar angles produced sex estimations with an accuracy of around 74%, for both 3D model and 2D photograph angles. As the latter are consistently around 1° smaller than the 3D model angles, the mean difference between male and female inclination angles is about the same for the two data sets, i.e. around 6° for glabellar angles (Table 2). It is therefore not surprising that discriminant functions based on the two data sets perform roughly on par, showing that both data sets are about equally useful for sex estimation.

Discriminant functions based on supraglabellar angles produced sex estimations with an accuracy of around 62% (Table 3). This lower performance is partially caused by a larger variation in the angles measured from supraglabella (Supplementary Table S2), arguably related to supraglabella being difficult to precisely locate on the skull, and partially caused by a smaller female/male difference in the glabellar angles (around 4° for supraglabellar angles derived from both 2D photographs and 3D models; Table 2). This smaller sex difference is probably related to the glabellar inclination angle not only reflecting the slope of the forehead, but also the prominence of glabella itself. Such a notion is supported by Spearman’s rank correlation coefficients of around 0.63 for glabella scores versus glabellar angles, while the correlation between glabella scores and supraglabellar angles is only around 0.4. The strong correlations between the forehead slope, the frontal eminences, and the glabella, led Rogers [10] to suggest that they should be considered a single point of evidence for sexing remains. However, while these features share a common downward and forward growth, related to the development of the frontal and nasomaxillary complexes [33], the presence in our sample of multiple crania with a clear frontal slope and a small glabella (e.g. Supplementary Fig. S3) suggest that these features are affected by different growth factors, and may not share the same timing of development and appearance.

The best sex estimation results (around 80% accuracy) were obtained based on trait scores of glabella from 1 to 5, evaluated from 3D models (Table 4). These good results are in line with previous research [10, 11, 34, 35], but the high frequency of interobserver disagreement in glabella scores (about 30%; Supplementary Tables S1 and S3) makes this method unsatisfactory for legal cases: it does not live up to the Daubert standard [19, 20]. It is possible that the glabellar scores would have been more consistent if scoring had been done from the physical crania, rather than from images. An opposite trend was observed in the frontal inclination scores: dividing the sample into two categories of “sloped” versus “vertical” frontal inclination yielded interobserver errors in the range 15-20%, and a sex determination accuracy of about 71% (Table 4; Supplementary Tables S1 and S3). This accuracy is only marginally lower than the 73% achieved by glabellar inclination angles, and much higher than the results of Williams and Rogers [10, 11] who scored frontal inclination from physical crania. We attribute this better performance to lower error rates in assessing angles from images presenting the cranium in perfect lateral profile rather than from real skulls, although population differences and not using an “indeterminate” group may also be contributing factors: it has been argued that fewer character traits should improve interobserver accordance [12], and using an indeterminate group affected the final accuracy in Rogers as well as Williams study [10, 11].

Sex estimation from angular measurements better classified females than males (Table 3), as female crania displayed a less variable inclination angle (Table 2), while estimations based on trait scores performed better for males in most cases (Table 4). The trait score results however vary between observers AP and SBS (Supplementary Tables S1 and S3), but the smaller variation in measured female inclination angles might have a physiological explanation: craniofacial growth in males does not cease until early adulthood, resulting in a longer period during which development disruptions or alterations can occur. Since female crania complete their growth much earlier than males [10], we would expect their frontal bones to exhibit less shape variation in adulthood.

In summary, this study presents the first test of skull dimorphism in a Croatian sample [24, 25, 36–38]. Although a clear sexual dimorphism in frontal inclination was observed, the sex estimation accuracy was not as good as that obtained by e.g. long bone measurements. Our results show that frontal inclination measurements may be a useful component in a multi-feature system for statistical sex determination of Croatian skeletons. A clear and systematic difference was observed between frontal inclination angles measured from 2D photographs and from screenshots of 3D models. Both types of measurements appear to be useful and can be recorded with low observer error, but angles obtained from these two different sources should not be combined. As there is no “known-sex” skeletal collection in Croatia, the study was conducted on an archaeological skeletal sample where sex had been determined from sub-cranial bones with an accuracy of around 95 to 98 percent, leaving a small uncertainty in the sex estimation results but not in the main methodological finding of this study, i.e. the systematic difference between the 2D and 3D angle measurements. Future research should investigate the frontal inclination angles for other populations, possibly for different age groups, and try to elucidate the factors involved in the growth and development of the frontal bone, as well as the factors responsible for the difference in angles derived from 2D and 3D images.

## Compliance with ethical standards

The 3D CT models were recorded in compliance with the current Croatian laws and ethical guidelines for the handling of human remains.

## Acknowledgments

We thank the Department of Diagnostic and Interventional Radiology at the Clinical Hospital Dubrava, Croatia, and in particular the Head of Dept. Prof. Boris Brkljacic, for facilitating CT scanning of the studied crania.

